# CarveMe-GutMicrobes: Automated Metabolic Model Reconstruction for Gut Microbial Species and Communities

**DOI:** 10.64898/2026.06.26.734454

**Authors:** Arianna Basile, Indra Roux, Aditi Madkaikar, Francisco Zorrilla, Stephan Kamrad, Kiran Raosaheb Patil

## Abstract

Genome-scale metabolic models (GSMMs) are important aids towards system-level understanding of the metabolic physiology of the gut microbes and for rational microbiome engineering. While large-scale repositories of GSMMs for gut-associated bacteria are available, strain-level variability and the continuous discovery of novel taxa through metagenomics and culturomics underscore the need for scalable, ab initio reconstruction tools. Here, we present CarveMe-GutMicrobes, a client-side framework for rapid reconstruction of metabolic models directly from (meta)genomic input. Building upon the original CarveMe framework, CarveMe-GutMicrobes incorporates an expanded, gut-microbe-centric biochemical database that includes reactions, metabolites, and gene-protein-reaction (GPR) associations curated specifically for Bacteria and Archaea inhabiting the human gut. The tool supports taxonomic restriction of the reference database to improve context-specific accuracy. CarveMe-GutMicrobes models demonstrated high predictive performance performance against gene essentiality and metabolite secretion datasets. By integrating curated resources, extending reaction coverage, and offering new empirical datasets, CarveMe-GutMicrobes provides a scalable platform for high-resolution metabolic reconstruction towards broader adoption of GSMMs in gut microbiome research.

## Introduction

The gut microbiota contributes essential protective, structural, and metabolic functions to host physiology and health (Colella *et al*, 2023). Genome-scale metabolic modeling has emerged as an effective tool to study the metabolic capabilities of individual gut microbes and the emergent properties of microbial communities (Agus *et al*, 2021; Esvap & Ulgen, 2021; Heinken *et al*, 2021; Colella *et al*, 2023; Quinn-Bohmann *et al*, 2025). Metabolic models enable the prediction of both competitive (Schäfer *et al*, 2023; Cerk *et al*, 2024) and cooperative (Blasche *et al*, 2021; Kost *et al*, 2023; Machado *et al*, 2021) ecological interactions, facilitate the study of strain-level functional divergence (Morlino *et al*, 2023; Keller *et al*, 2022), and allow for in-silico simulation of dietary interventions (Basile *et al*, 2020; Weston & Thiele, 2023) and pathogen invasion scenarios (Raghunathan *et al*, 2009; Catlett *et al*, 2020).

While bacteria constitute the dominant fraction of the human gut microbiota, the community also encompasses members of the Archaea domain, which remain relatively understudied despite their functional significance (Gaci *et al*, 2014; Fumagalli *et al*, 2025). Archaeal taxa, particularly methanogenic species, contribute to microbial metabolic networks through cross-feeding and redox balancing and can impact human health. For example, a metabolic synergy has been described between *Bacteroides thetaiotaomicron* and *Methanobrevibacter smithii*, in which cross-feeding of fermentation byproducts promotes mutual growth and reflects a syntrophic relationship within the gut microbiota (Catlett *et al*, 2022). Conversely, elevated levels of archaeal methanogens and their metabolic product, methane, have been associated with gastrointestinal disorders, including Crohn’s disease, suggesting a broader clinical relevance beyond benign symptoms like bloating (Houshyar *et al*, 2021; Basile *et al*, 2023).

Both bacterial and archaeal members influence host physiology and interspecies interactions via the secretion of small molecules. Gut-derived metabolites have critical immunomodulatory effects on host tissues. For instance, bacterial catabolites of tryptophan—such as tryptamine, indole, and indole-3-propionic acid (IPA)—alongside short-chain fatty acids (SCFAs), influence epithelial barrier integrity and modulate host immune responses, including anti-tumour immunity and responses to immunotherapy (Hezaveh *et al*, 2022; Xue *et al*, 2023). Understanding how production of these and other gut bacterial metabolites is modulated by nutrient availability and community context represents a critical step toward mechanistic modelling of microbiome function. Constraint-based metabolic models offer a systematic framework to explore environmental and genetic determinants of metabolite secretion and thereby to generate testable hypotheses for decoding and modulating microbe-microbe and microbiome-host interactions.

Metagenomics is the most used approach for characterising gut microbiota due to its scalability and culture-independence (Joseph & Pe’er, 2021). Genome-scale metabolic models (GSMMs) (Zampieri *et al*, 2023; Luo *et al*, 2023) can be built directly from the species-level genomes, metagenomics data (Zorrilla *et al*, 2021), or from metagenome assembled genomes (MAGs) (Almeida *et al*, 2019; Nayfach *et al*, 2019; Palù *et al*, 2022). The GSMMs are often applied using the constraint-based reconstruction and analysis (COBRA) family of methods such as Flux Balance Analysis (FBA) to simulate, e.g., minimal media requirements (Gelbach *et al*, 2024), gene essentiality (Kaynar *et al*, 2023), metabolite secretion (Quinn-Bohmann *et al*, 2024), cross-fed metabolites (Gabrielli *et al*, 2023; Basile *et al*, 2020) and the degree of competition between community members (Machado *et al*, 2021; Schäfer *et al*, 2023).The fidelity of these simulations is directly contingent on the quality of the underlying models, particularly the accuracy of gene annotations and the representation of gene-protein-reaction (GPR) associations. To support this, manually curated resources such as the BiGG Models database (Norsigian *et al*, 2020) and the AGORA2 collection (Heinken *et al*, 2023) stands out as two extensive GSMM collections. These repositories provide models with rigorously curated reactions, each bearing unique identifiers and comprehensive GPR rules encoded using Boolean logic to represent enzyme isoforms and multimeric complexes (Filippo *et al*, 2021). These rules describe the relationship between gene products, such as enzyme isoforms or subunits, involved in catalysing a particular reaction, using Boolean logic. The quality of the GPR rules incorporated into GSMMs heavily influences the reliability of formulated hypotheses (Zampieri *et al*, 2023). Despite their breadth and depth, both AGORA2 and BiGG are semi-static resources built from isolated genomes, making them suboptimal for dynamic applications such as community modelling involving novel taxa or for benchmarking emerging model-building methodologies. To address this, several automated tools have been developed for de novo genome-scale metabolic model reconstruction from genomic data, including CarveMe (Machado *et al*, 2018), gapseq (Zimmermann *et al*, 2021), ModelSEED-based pipelines (Heinken *et al*, 2021), Pathway Tools (Karp *et al*, 2010), RAVEN (Wang *et al*, 2018), and Merlin (Capela *et al*, 2022), each relying on different databases and reconstruction strategies.

A distinguishing feature of CarveMe, which we contributed to the development of, is the top-down reconstruction strategy grounded in the use of a ‘universal model’ of microbial metabolism. The universal model is meant to be a comprehensive representation of biochemical reactions within the group of target organisms (e.g., bacteria) and is curated as a whole to improve accuracy regarding reaction directionalities and metabolic connectivity, amongst others. The reconstruction process for a given species or a strain begins with the universal model, reducing it systematically to minimise the number of reactions with no genomic evidence, while retaining the feasibility of the model for biomass formation and, if specified, maintaining compatibility with growth on a specific nutrient mixture. The curation of the universal model and the maintenance of the model feasibility for simulations together make CarveMe applicable to a variety of large-scale metabolic modelling problems (Blasche *et al*, 2021; Schäfer *et al*, 2023), which can be otherwise difficult to tackle due to the need for extensive manual curation and/or computationally costly gap filling process. Recent benchmarking studies have shown that CarveMe performs competitively with state-of-the-art approaches, providing high-quality draft models with good accuracy and scalability, and therefore is used here as reference.

The first CarveMe release used the BiGG database as the foundation for its universal model primarily focussed on bacterial metabolism. Since, several developments have been made, experimentally and computationally, towards enhancing the knowledge of bacterial metabolism, and especially that for gut bacteria. AGORA2 comprises a vast repertoire of GPR rules, and their integration in the CarveMe universe can be highly advantageous for the COBRA scientific community. However, challenges remain for full incorporation of AGORA2 into the *CarveMe* framework, notably the differing nomenclatures that hinder compatibility and cross-simulation between AGORA2- and BiGG-based reconstructions.

Here we present CarveMe-GutMicrobes, a gut microbiota focussed release of CarveMe comprehensively representing the metabolism of anaerobic microbes. The main features include the integration of the GPR rules from the AGORA2 collection and relevant databases such as the Transporter Classification Database (TCDB) (Saier *et al*, 2021) and UniProt (UniPac 90) (UniProt Consortium, 2023). Compared to its initial release, the CarveMe-GutMicrobes universal database is substantially expanded with 1,618 new reactions and nearly 70,000 new GPR rules. Further, an additional feature is included enabling taxonomic subsetting of the repertoire to match the target organism and thereby reduce the false positive matches.

A major hurdle in developing and applying metabolic models to gut bacteria is the lack of systematic experimental data, especially for gene essentiality, which has been a foundation of GSMMs development. To partly address this gap, we compiled a new dataset of gene essentiality profiles for three non-model gut bacterial species, which we used to test and evaluate the predictive performance of our metabolic models. Demonstrating their utility, the models recapitulate, with good accuracy, gene essentiality and metabolite production.

## Materials and methods

A dedicated GitHub repository (https://github.com/arianccbasile/carvemegut) provides all the needed scripts to build the three elements constituting the new universal model: the universal bacterial and archaeal reconstructions, the universal protein database, and the GPR compendium, as well as the pipeline for the reconstruction of metabolic models. The software is also accessible via the Python Package Index (PyPI) and can be installed using pip. A separate dedicated GitHub repository (https://github.com/arianccbasile/CarveMe-GutMicrobes_paper) provides hands-on tutorials and best-practice guidelines for using CarveMeGut-Microbes.

### Universal bacterial model

The draft model was built integrating the reactions from the 7,302 GSMMs of the AGORA2 collection (Heinken *et al*, 2023) with the latest BiGG universe (version 1.6; http://bigg.ucsd.edu/) (Norsigian *et al*, 2020). Following previously defined standard procedures (Thiele & Palsson, 2010), the universe model was refined implementing key curation steps. Reactions of biomass and ATP maintenance were added; the hydrogen stoichiometry was fixed as well as protons and charges balance. The metabolite repertoire was curated filling the missing formulas, and creating sink and exchange reactions. Additionally, blocked reactions and dead-end metabolites were identified through flux variability analysis and removed from the model. The steps listed above are included in the sanity check performed by the function “curate_agora” available in the Github CarveMe-GutMicrobes_paper repository. All available metadata are saved as annotations in the resulting SBML. A set of reactions were manually added to the resulting model using the function add_reaction_from_string in the reframed (https://github.com/cdanielmachado/reframed/tree/master) python module.

### Specialised taxonomic templates

Following established protocols for top-down metabolic model reconstruction (Seaver *et al*, 2021; Machado *et al*, 2018; Zimmermann *et al*, 2021), we generated tailored metabolic universes for gram-positive, and gram-negative bacteria and Archaea. The gram-positive template includes characteristic biomass components such as glycerol teichoic acids, lipoteichoic acids, and a peptidoglycan unit, representing the thick cell wall typical of this group. The gram-negative template incorporates phosphatidylethanolamines, murein, and a lipopolysaccharide unit, reflecting the double-membrane architecture and outer membrane components. For Archaea, we accounted for several distinct features, including ether-linked lipids in the cell membrane, unique isoprenoid-derived lipid backbones, and the absence of peptidoglycan, with pseudomurein or S-layer proteins typically forming the cell wall instead. Additionally, we incorporated pathways specific to archaeal metabolism, with a particular focus on methanogenesis. The template covers the three major types of methanogenesis: hydrogenotrophic, methylotrophic, and acetoclastic, reflecting the diverse range of substrates (H_2_/CO_2_, methylated compounds, and acetate, respectively) that can be converted to methane.

### Protein database curation

Together with the amino acid sequences of all genes in the BiGG database, we utilized the genome sequences of 6,223 organisms included in the AGORA2 collection (version 2.01), as accessed from the NCBI database (Sayers *et al*, 2022). To represent the archaeal community, 1,167 MAGs of Archaea from the work of Chibani and colleagues were used (Chibani *et al*, 2022). The obtained genomic sequences were translated into proteins using prodigal (2.6.3-gcc-5.4.0-lftcuch) (Hyatt *et al*, 2010). Additionally, 25,897 protein sequences from UniRef90 and 21,362 from TCDB and other transporters from UniRef90 were downloaded from gapseq’s GitHub repository (Zimmermann *et al*, 2021). Transporters from the gapseq repertoire (Zimmermann *et al*, 2021) were clustered using cd-hit (parameters: -c 0.9, -n 5) (Fu *et al*, 2012). After the clustering, 20,960 unique sequences were left. The whole repertoire of sequences obtained from UniRef90 and TCDB is stored as a single protein FASTA file, most of the ids were edited and special characters removed. A total of 443,067 protein sequences were obtained from AGORA2. The protein sequences extracted from AGORA2 are stored in the directory called proteins_divided.

### Taxonomical carving of the protein database

The parameters --taxonomical_level (-tl) and --taxonomy (-t) are introduced. Knowledge on the taxonomy of the organism of interest can be used to refine the protein database which is incorporated as part of the reference for drafting metabolic models. Based on user input, specific sequences are filtered from the collection of all the sequences sourced from the AGORA2 repertoire. Accordingly, the final protein file will contain the selected sequences as well as all the sequences from TCDB, UniRef90, BiGG.

The function makedb of DIAMOND (Buchfink *et al*, 2015) was used to create a database from the final protein file. When running CarveMe-GutMicrobes, the user-provided input genomes are aligned to the resulting database in order to identify the corresponding homologous sequences in the protein database.

### Gene-protein-reaction compendium

All the sequences extracted from AGORA2 and the collection of Chibani *et al* are annotated with gapseq (function find) and the corresponding BiGG reactions are obtained through linking the ModelSEED ids with the corresponding BiGG ids. All the transporters from Chibani and AGORA2 datasets are annotated with gapseq (function find-transport) (Zimmermann *et al*, 2021), and the ids are converted from ModelSEED (Seaver *et al*, 2021) nomenclature to BiGG nomenclature (Norsigian *et al*, 2020). Original AGORA2 GPRs are extracted using in-house developed scripts and linked to the corresponding sequences. Extraction of the GPRs followed multiple quality checks: 1) the gene id of each gene was cross-checked between the GSMM and the corresponding genome; 2) Eggnog annotation was performed for all the protein sequences; 3) Eggnog annotation was cross-checked with the annotation obtained from AGORA2; 4) when the annotations were not consistent, additional manual checks were performed including checking the gapseq annotation.

Finally, the GPRs are stored in a single file in the subfolder data/generated. After mapping the genomes of interest on the protein database, gene scores are converted to protein scores by selecting the lowest score among all the subunits that make up a protein complex. Subsequently, protein scores are transformed into reaction scores by adding the scores of all isozymes that can facilitate a particular reaction. Throughout this process, custom GPR associations are created for the organism being reconstructed.

### Implementation

CarveMe-GutMicrobes is implemented in Python 3.8. It requires the reframed python package (version 1.2.1) for metabolic modelling, which provides an interface to common solvers (Gurobi, CPLEX), and import/export of SBML files through the libSBML API (46). In this work, all MILP problems were solved using the IBM ILOG CPLEX Optimizer (version 22.1.0.0). Furthermore, gurobi can also be used as a solver.

### Model testing and validation

The newly developed CarveMe-GutMicrobes reconstruction workflow was rigorously assessed for effectiveness in replicating metabolite production across five microbial strains, as well as gene essentiality data. The criteria utilised to evaluate the simulations results obtained by comparable softwares are as follow:

Precision: TP / (TP + FP)

Sensitivity (also known as Recall): TP / (TP + FN)

Specificity: TN / (TN + FP)

Accuracy: (TP + TN) / (TP + FP + TN + FN)

F1-score: 2 * (Precision * Sensitivity) / (Precision + Sensitivity)

Matthew correlation coefficient: (TP∗TN- FP∗FN)/(sqrt((TP+FP)∗(TP+FN)∗(TN+FP)∗(TN+FN)) (Chicco and Jurman, 2020)

The Matthews correlation coefficient (MCC) yields high scores exclusively when predictions demonstrate success across all four categories of the confusion matrix (true positives, false negatives, true negatives, and false positives).

Where:

TP = True Positive (correctly predicted positive samples)

TN = True Negative (correctly predicted negative samples)

FP = False Positive (incorrectly predicted positive samples)

FN = False Negative (incorrectly predicted negative samples)

### Prediction of secondary metabolites production

To assess the effectiveness of our approach in predicting bacterial metabolism under anaerobic conditions and to compare it to other existing model reconstruction tools, growth of bacteria in anoxic environments was simulated and the production rates of a range of secondary metabolites derived from the metabolism of amino acids was assessed. To ensure consistency, all the compared species models were reconstructed and gap-filled using CarveMe, and CarveMe-GutMicrobes. Additionally, when available metabolic reconstructions of the AGORA2 collection representing the organisms of interest were obtained from the Virtual Metabolic Human platform (VMH; https://www.vmh.life/) (Noronha *et al*, 2019). All the simulations were performed using reframed (https://github.com/cdanielmachado/reframed).

The function load_cbmodel was used to import models, *Environment.complete* was used to constrain the reconstruction to a rich nutrient environment. Oxygen uptake was set to 0 using the BiGG or ModelSEED nomenclature as required. This medium was used to constrain the model using *Environment.from_compounds*. Flux-Variability-Analysis (FVA) was used to estimate the maximum possible release of a range of secondary metabolites. The simulations were obtained using the function *FVA* of reframed. The metabolites were considered as 1 (present) or 0 (absent) if their production was over or below the average of the negative controls for that metabolite. More details on the experimental framework are reported below. IBM ILOG CPLEX optimizer was chosen as linear programming solver.

### Gene essentiality benchmark

To assess the effectiveness of CarveMe-GutMicrobes, genome-scale metabolic models were used to predict gene essentiality. In detail, using the GPR mappings we performed in silico single gene deletion analysis. The predicted growth/no-growth phenotypes from the network models were compared to experimentally determined growth phenotypes of gene deletion mutants using contingency tables. The GSMMs of the three organisms considered for this benchmark i.e., *Bacteroides thetaiotaomicon*, *Parabacteroides merdae* and *Bifidobacterium breve,* are drafted using CarveMe and CarveMe-GutMicrobes and the AGORA2 reconstructions of *Bacteroides thetaiotaomicron* (VPI_5482), *Parabacteroides merdae* (ATCC_43184), and *Bifidobacterium breve* (UCC2003_NCIMB8807) are obtained from VMH (Noronha *et al*, 2019).

Adopting the same assumptions as Machado and colleagues (Machado *et al*, 2018), genes were considered conditionally essential in the given growth environment if the predicted growth rates were below 0.01 h-1. To enable coherent comparison, only genes present in the GSMMs of the corresponding species from all three softwares were considered. To ensure consistency across different genome versions used in model reconstruction, gene names were standardised based on sequence comparisons using diamond (v. 2.0.15). In all the cases, the simulations were conducted using the flux variability analysis algorithm and under a rich nutrient environment lacking oxygen.

For *B. thetaiotaomicron*, the list of essential genes was obtained intersecting the lists of essential genes found in two independent works (Goodman *et al*, 2009; Liu *et al*, 2021). The gene essentiality data for P. merdae was derived from a dense transposon mutant library of P. merdae ATCC 43184 selected in solid mGAM media (Voogdt *et al*., 2025). TnSeq reads were mapped to the annotated *P. merdae* genome GenBank: CP102286.1, and gene essentiality predicted using TRANSIT V.3.2.7 with analysis with Hidden Markov Model (HMM) trimming at both 10% of C- and N- gene terminal (DeJesus *et al*, 2015). The essential genes specific to *B. breve* were obtained from Ruiz and colleagues (Ruiz *et al*, 2017).

### Archaea benchmark

Five different Archaea species (*Methanobrevibacter ruminantium, Methanomassiliicoccus luminyensis, Methanosphaera stadtmanae, Sulfolobus solfataricus, Methanobrevibacter smithii*) spanning three different genera were benchmarked. Using corresponding genomes from the RefSeq database (O’Leary *et al*, 2016), the models were reconstructed with CarveMe and CarveMe-GutMicrobes softwares. The reframed framework was used to load the models and simulate a nutrient rich complete environment. The oxygen uptake was set to 0 to emulate an anaerobic environment, and the resulting medium was used to constrain the models. FBA simulations were used to simulate metabolite uptake and secretion profile in a nutrient rich anaerobic environment. This analysis was carried out for the GSSMs of the five archaeal species obtained from AGORA2 collection as well as ones reconstructed using CarveMe and CarveMe-GutMicrobes. The secreted or uptaken metabolites were considered present (1) or absent (0) and were then compared, using the benchmarking criteria described above, with the records extracted from the NJC19 database (Lim *et al*, 2020).

### Metabolic reconstruction of HumGut MAGs

Metagenome-assembled genomes (MAGs) from the HumGut collection (Hiseni *et al*., 2021) were downloaded and used as input for genome-scale metabolic model reconstruction. For each MAG, metabolic models were generated using CarveMe and CarveMe-GutMicrobes. The resulting genome-scale metabolic models were subsequently employed for downstream metabolic analyses. To compare the diversity of metabolic models generated by the two methods (CarveMe and CarveMeGut), a binary matrix of reaction presence/absence was constructed for selected GSMMs representing strains of the same species. In this matrix, rows represent strains, columns represent reactions, and binary values indicate the presence (1) or absence (0) of each reaction in the model. For each method, the Jaccard distance was calculated between all pairs of models, resulting in a pairwise distance matrix. Statistical significance of the differences between CarveMe and CarveMe-GutMicrobes distances was assessed using a paired Wilcoxon test.

## Results and discussion

The CarveMe universe comprises three interconnected components: the protein database, the universal metabolic model, and the GPR compendium. *CarveMe-GutMicrobes* builds upon an expansion and refinement of all three elements. We initiated this process by constructing a new universal bacterial metabolic model through integration of the BiGG database (version 1.6) (version 1.6) (Norsigian *et al*, 2020) with the AGORA2 collection. The BiGG database encompasses 108 genome-scale metabolic reconstructions spanning 48 different species, while the AGORA2 collection, at the time of our access, contained 7,302 metabolic reconstructions. All of these were used in the creation of the universal reconstruction at reaction level; of which 6,223 could be added to the final protein database and to the GPR compendium following quality control (Methods). A total of 1,167 archaeal genomes from the most updated collection of gut archaeal MAGs (Chibani *et al*, 2022) were also integrated in the protein database and in the GPR compendium. Figure 1 summarizes the overall study design and lists the databases used for benchmarking.

**Figure 1:**
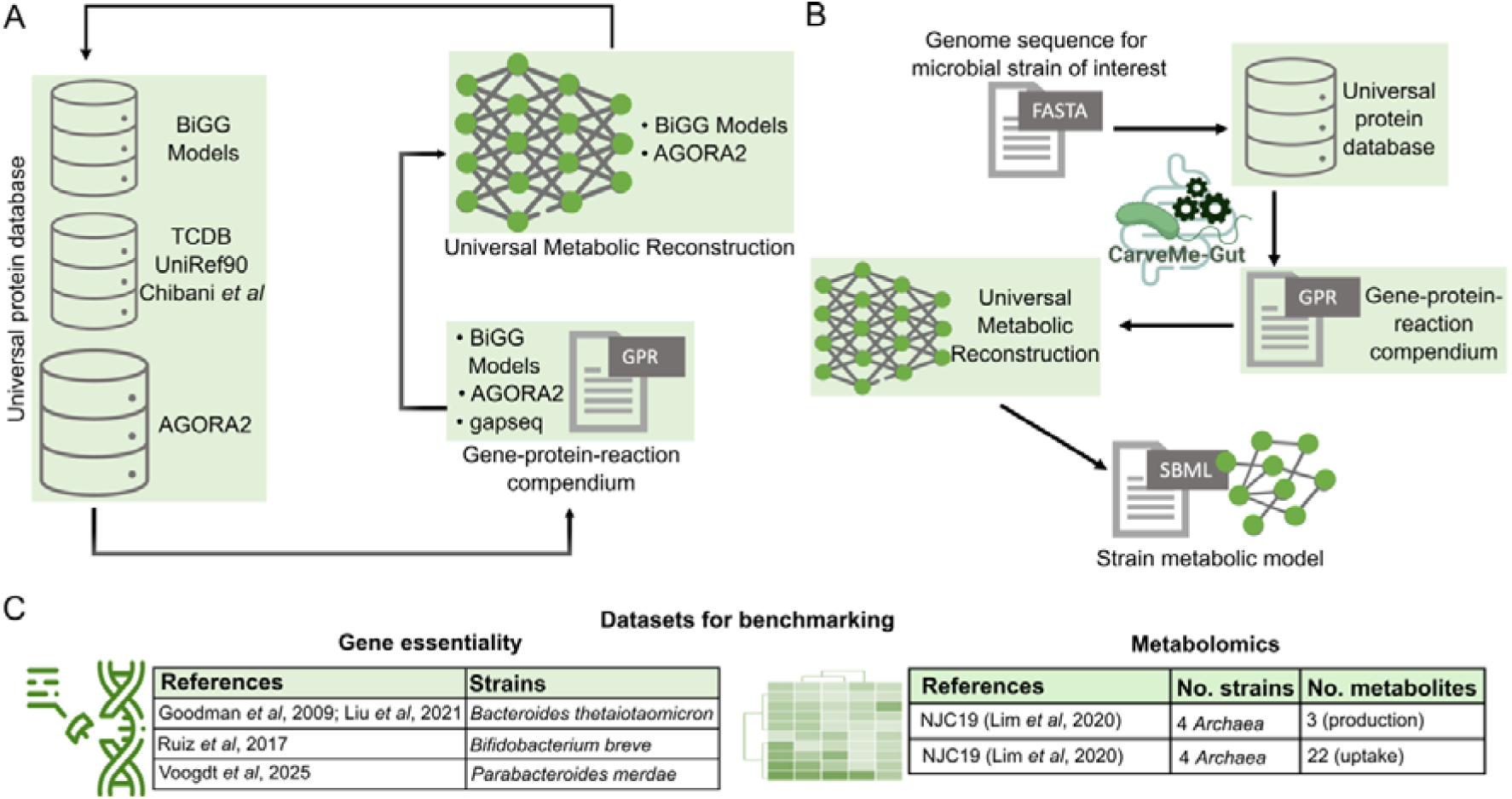
(A) Overview of the extensions and updates made for the CarveMe-GutMicrobes release. The universal protein database was expanded with sequences from AGORA2, TCDB, UniRef90 as well as the MAGs collection from Chibani et al. (Chibani *et al*, 2022); the GPR compendium was expanded with rules from gapseq and AGORA2; the universal metabolic reconstruction was created by integrating reactions from BiGG models database and AGORA2. (B) Overview of the reconstruction process using CarveMe-GutMicrobes. The workflow starts with the input fasta file and produces a simulation-ready GSMM. The genome of interest (as nucleotide or protein fasta) is imputed in the CarveMe-GutMicrobes software and aligned on the target database which can be refined taxonomically (facultative). Following the annotation, relevant gene-protein-reaction relationships are obtained and, accordingly, relevant reactions are carved from the universal SBML template in order to obtain the final model. (C) Overview of the datasets used for benchmarking the CarveMe-GutMicrobes.

### Construction of an Integrated Universal Metabolic Reconstruction

The bacterial universal reconstruction was built by starting with the BiGG bacterial universe (v.1.6) and integrating reactions from AGORA2. Then, the model underwent a series of curation processes, as outlined in the Methods section. These steps included, among others, reaction directionality assessment, validation of elemental balance, removal of energy-producing cycles, and incorporation of a template biomass composition. The final bacterial model includes three compartments (cytosol, periplasm, extracellular space), features 3,030 metabolites (representing 1,778 distinct compounds) (Table S1) and 6,001 reactions (3,721 enzymes, 1,618 transporter-mediated exchanges, and 662 inter-compartment metabolite exchanges). For a more reliable depiction of fatty acid synthesis and glycerophospholipid metabolism pathways, we added the additional biosynthetic pathways of Palmitoyl-ACP (n-C16:0ACP), Myristoyl-ACP (n-C14:0ACP), Octadecanoyl-ACP (n-C18:0ACP), Acetoacetyl-ACP, Cis-octadec-11-enoyl-[acyl-carrier protein] (n-C18:1) were added for a more reliable depiction of fatty acid synthesis and glycerophospholipid metabolism pathways. Accordingly, lipid metabolism is the metabolic sub-system with the highest number of newly included reactions (Figure 2A).

**Figure 2:**
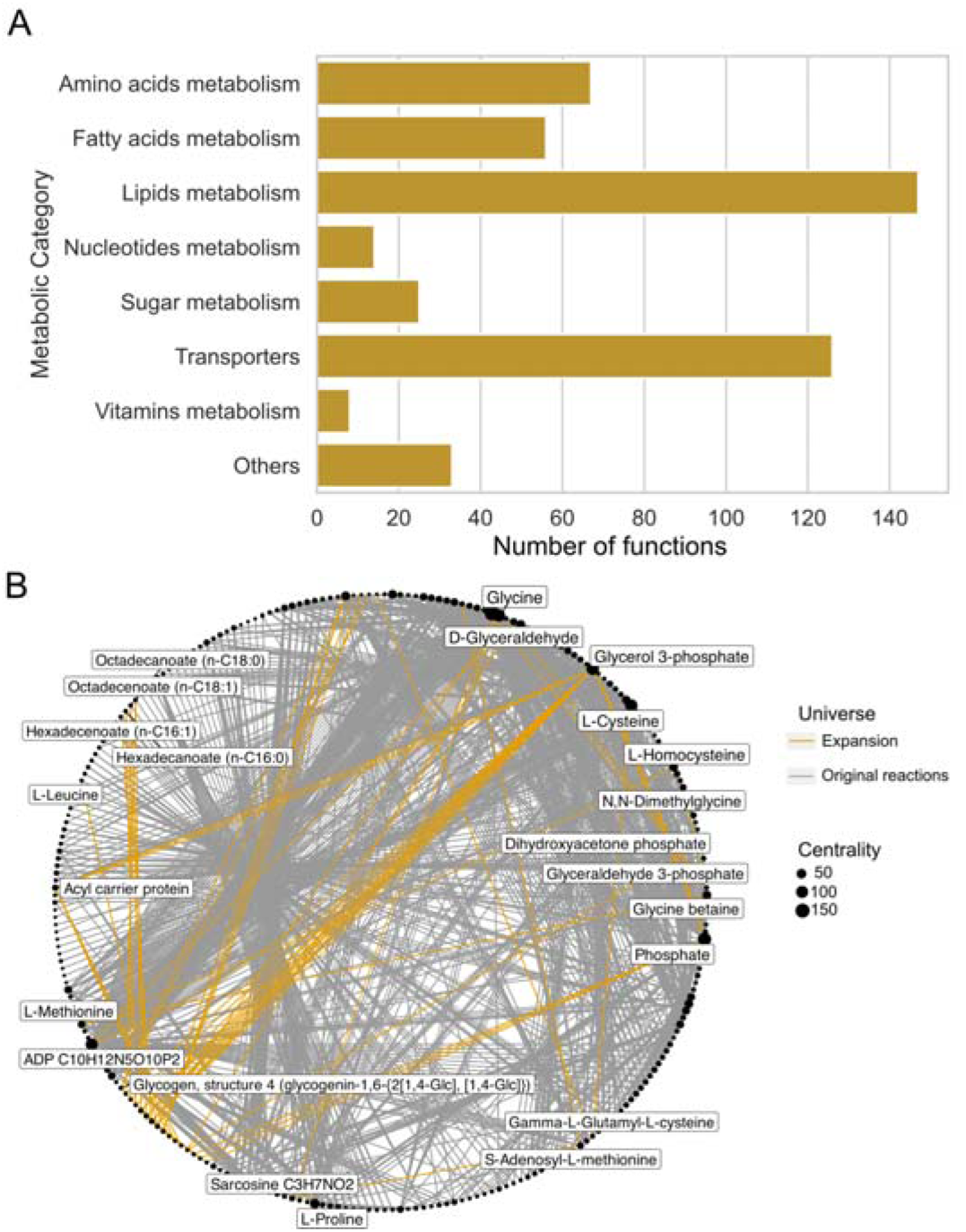
(A) Distribution of the newly added reactions in the expanded universe of CarveMe-GutMicrobes across different metabolic pathways; (B) Overview of the expanded universe highlighting new reactions in dark yellow. Newly added reactions involving Cysteine, Glycine and Proline are marked by naming the key substrate nodes. Reactions for the other classes of amino acids are summarised in Figure S1.

Further, in the integrated universal metabolic reconstruction the reaction repertoire of L-Phenylalanine, L-Histidine, and L-Tyrosine showed an increase from 18 to 23 (28%), from 18 to 25 (39%), and from 21 to 31 (48%), respectively (Figure S1A, S1B). Among the uncharged amino acids, the selenoamino acid metabolism involving L-serine, was added (Figure S1C). Another group of newly added reactions is associated with the metabolism of sarcosine (Figure 2B), a key intermediate in glycine metabolism.

Secondary metabolites play a crucial role in shaping the interactions between gut-microbes and the host (Tintelnot *et al*, 2023; Visconti *et al*, 2019). To increase the coverage of secondary metabolites in metabolic models, we added several secondary metabolism pathways (Methods). For example, reactions for synthesis of histamine, a biogenic amine, important for regulation of vascular permeability and promotion of mucus secretion (Mou *et al*, 2021), were included. The enzyme histidine decarboxylase (K01590) is responsible for producing histamine through decarboxylation of the amino acid histidine. In the new reconstruction, this reaction is integrated with two different IDs, HDC and HISDC. The main difference lies in their proton balance: HDC includes two protons (2.0 h_c) as reactant, while HISDC only includes one proton (1.0 h_c) and the type of cofactor used. Moreover, synthesis of tyramine from tyrosine by L-Tyrosine carboxy-lyase (K01592) was included as TYRCBOX (Figure S1C). A full list, complete of metadata, of all the new reactions and metabolites is available in TableS1. In addition to the universal reconstruction, we also developed specialised templates for Archaea, and for gram-positive and gram-negative bacteria. It is important to note, however, that mobile genetic elements such as prophages are only partially represented and not explicitly distinguished in the GPR framework. While prophages typically contribute limited core metabolic functions, they can encode auxiliary enzymes, transporters, or regulators that influence strain-specific metabolism, and their inclusion could enhance the resolution of strain-level metabolic modelling. By selecting the template most appropriate for a given organism, users can generate models that better reflect group-specific physiology and metabolic capabilities, while acknowledging that certain mobile genetic contributions remain underrepresented.

### The Integrated protein database and GPR compendium

#### Transporters and enzymes databases

The protein database and GPR compendium was extended using TCDB and UniRef90 to improve the coverage of transporters. We started by downloading 7,390 genomes of Archaea and Bacteria from European Bioinformatics Institute repository (ftp.ebi) and NCBI; these included all the MAGs from (Chibani *et al*, 2022) as well as taxonomically heterogeneous isolate genomes included in the AGORA2 collection (Heinken *et al*, 2023). The genomes were then annotated using gapseq (Zimmermann *et al*, 2021) which uses TCDB and part of UniRef90 for transporters annotation, to identify and annotate the transporters, and the nomenclature of metabolites was translated from ModelSeed to BiGG. To extend the database with non-transporter metabolic enzymatic reactions and sequences, we selected a subset of 286 microbial genomes covering taxonomically heterogeneous species for gapseq reactions annotations (using UniRef90for enzyme annotation). The analysis resulted in a total of 21,473 protein sequences forming 60,456 GPRs (Figure 3A). These GPRs cover diverse metabolic pathways, among which the most represented (18 %) is amino acid metabolism (Figure 3B).

**Figure 3:**
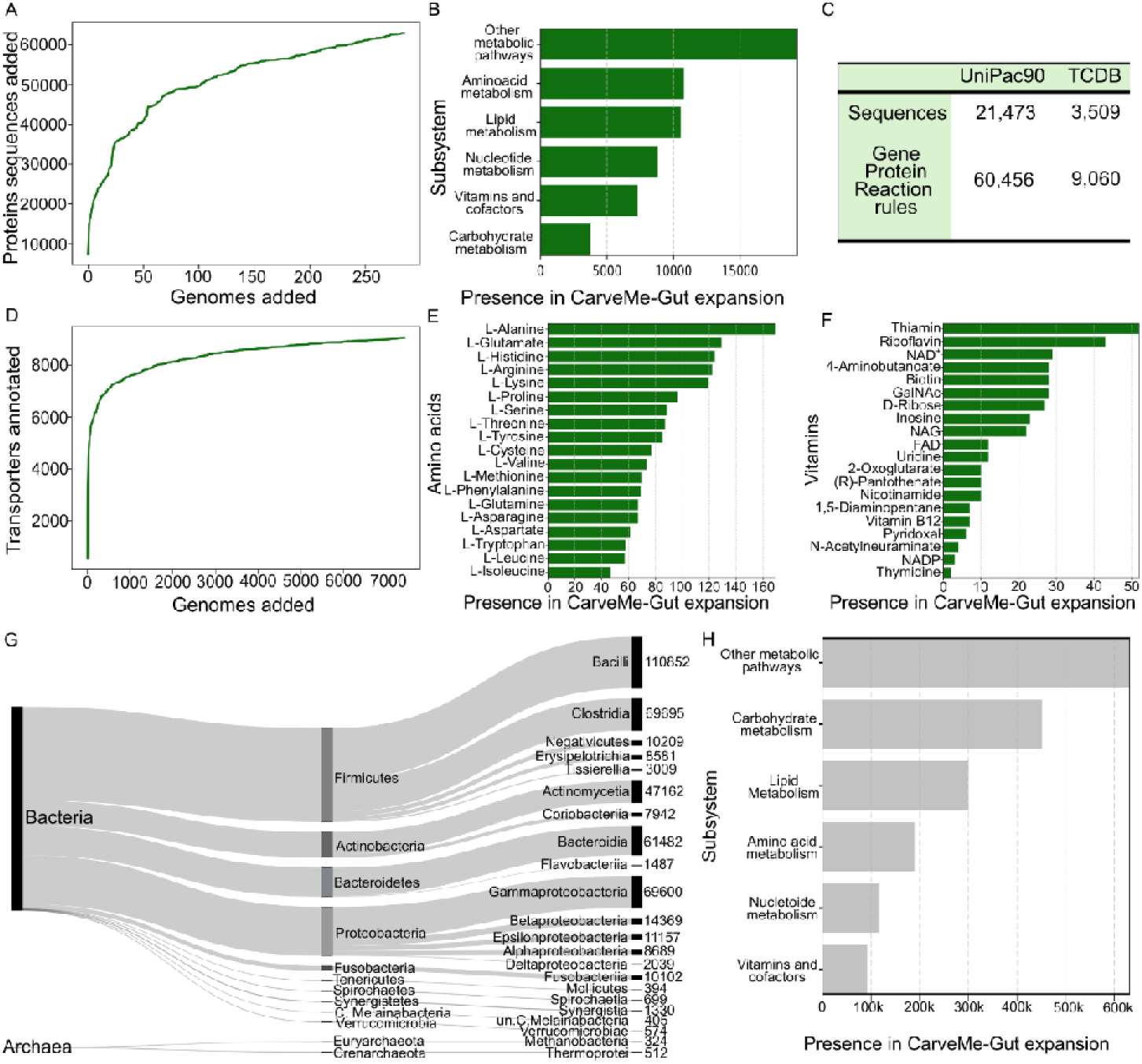
Overview of the sequences and GPRs added using UniRef90, TCDB, and AGORA2. (A) Protein sequences per genome added as reactions; (B) Distribution of GPRs added using the UniRef90 across custom-defined metabolism categories; (C) Number of sequences and GPRs added from UniRef90 and TCDB; (D) Increase in the sequences annotated as transporters as function of added genomes; (E) Distribution of transporters of added from TCDB across different amino acids; (F) Distribution of of vitamin transporters added from TCDB; (G) Taxonomic coverage of the extracted AGORA2 database of sequences; (H) GPRs added in the AGORA2 expansion based on custom defined supra metabolism categories.

The total number of sequences mapping to transporters is 3,509 spanning 9,060 gene protein reaction rules (Figure 3C). The transporters link to a total of 179 unique metabolites including amino acids (Figure 3D) and vitamins (Figure 3E). Consistent with the known transporter biochemistry, many amino acid transporters are non-specific, transporting a range of amino acids with similar properties, while vitamins transporters are generally specific (Celis and Relman, 2020).

The gene annotation can be difficult for no-model organisms and are constantly expanding as functional genetics is being developed. To ensure the users retain the flexibility, CarveMe-GutMicrobes reconstruction tool provides an option to use Eggnog (Huerta-Cepas *et al*, 2019) annotation using the “-egg” parameter, which takes as input the output from Eggnog (‘*.emapper.annotation’).

#### Further extension with reactions from AGORA2

Next, we extended the protein database as well as the GPR compendium with the GPRs from 7,302 genomes corresponding to the GSMMs of the AGORA2 collection (Heinken *et al*, 2023). GPRs from AGORA2 GSMMs were recursively extracted. Extraction of the GPRs was not feasible for all models/genes. This was due to multiple quality issues including inconsistent gene nomenclature and failed annotation cross-check with Eggnog (Huerta-Cepas *et al*, 2019).

In total, 440,613 protein sequences were extracted from 6,223 genomes and converted to the corresponding amino acid sequence using prodigal (2.6.3-gcc-5.4.0-lftcuch) (Hyatt *et al*, 2010). These sequences map to a total of 4,166,425 GPRs (Figure 3G, 3H). Extending the database with sequences and GPRs from AGORA2, UniRef90 and TCDB, as well as BiGG models, underpin this rise in sequences and corresponding GPRs. While the expansion likely increases the number of true positive reactions, it may also inflate the number of false positive reactions. To improve accuracy, i.e. reducing false positives but preserving true positives, the parameters -t and -tl are introduced for taxonomic pruning

Taxonomic pruning of the biochemical database

Two new parameters -tl (--taxonomic-level) and -t (--taxonomy) are introduced to accurately subselect the database according to evidence on the taxonomic annotation. The user can also decide not to specify the database of choice. Here are some examples on usage of -tl and -t:

- -tl Kingdom -t Archaea
- -tl Genus -t Alistipes
- -tl pHyLuM -t FiRmiCUTes

The two parameters must be used in tandem to correctly subset the sequence database. By adjusting both the taxonomic target and the level of taxonomic pruning, users can control the specificity and breadth of the sequences included. This ensures that relevant orthologues from closely related taxa are not inadvertently excluded while still focusing the database on the organism of interest. Careful tuning of these parameters allows for a balance between model accuracy and annotation coverage.

### Expanded Archaea template

The CarveMe Archaea template contains ether lipids instead of peptidoglycans in the composition of the cell membrane, but the specific GPR rules were stemming only from one species metabolic reconstruction, viz., *Methanosarcina barkeri* str. Fusaro model called iAF692. To make the Archaea template more comprehensive, 1,167 Archaea metagenome assembled genomes from the work of Chibani and colleagues (Chibani *et al*, 2022) as well as 7 isolate genomes from AGORA2 belonging to Euryarchaeota, Crenarchaeota, and Thermoplasmatota phyla were downloaded. The sequences were annotated using UniRef90 and TCDB as reference databases. In addition, the corresponding AGORA2 GSMMs were obtained to fetch Archaea-specific reactions for the specialised metabolic reconstruction Archaea template. This template model then underwent a curation process as described in the methods section. The new Archaea template includes 8,879 reactions and 3,972 metabolites, outnumbering the original template of 511 reactions and 206 metabolites.

### Model evaluation using experimental data

The CarveMe-GutMicrobes database and model reconstruction workflow was evaluated against multiple experimental datasets: (a) gene essentiality data for three species, among which *P. merdae* represents the most recent non-model gut bacterium with such data available, and (b) the production and uptake of key metabolites by archaeal taxa.

#### Bacteria

##### Gene essentiality analysis

We compared the AGORA2, CarveMe and CarveMe-GutMicrobes models for gene essentiality predictions in three species, viz., *Bacteroides thetaiotaomicron*, *Bifidobacterium breve*, and *Parabacteroides merdae*. Published gene essentiality data for the first two organisms are used for gene essentiality benchmarking, while in-house dataset was used for *P. merdae* (Voogdt *et al*., 2025).

The models from the three sources are generally comparable in terms of the number of reactions and genes with CarveMe-GutMicrobes featuring more reactions with GPRs than CarveMe (Figure 4A). Notably, AGORA2 includes a higher number of pathways, reflecting its manually curated reconstruction strategy and broader metabolic coverage, whereas CarveMe-derived models rely on automated, genome-driven reconstruction and therefore adopt a more conservative pathway assignment. For each species, the essentiality predictions were compared for the genes present in all the three models. Compared to CarveMe, CarveMe-GutMicrobes results feature higher TP and TN, as well as lower FP and FN. The performances of AGORA2 reconstructions are overall better than the CarveMe models. For *P. merdae* the results of AGORA2 metabolic reconstruction show less FN but a higher number of FP compared with the GSMM obtained with CarveMe-GutMicrobes. The sensitivity obtained using AGORA2 *P. merdae* is 0.48, higher than the one obtained with CarveMe-GutMicrobes. However, CarveMe-GutMicrobes results show higher precision, specificity, accuracy, F1-score and MCC-score than the corresponding AGORA2 GSMMs (Figure 4B-D). For *B. thetaiotaomicron*, a similar pattern emerges (Figure 4B, Table S3), with both CarveMe-GutMicrobes and AGORA2 models outperforming CarveMe across all metrics. Unlike *P. merdae*, the sensitivity scores of AGORA2 and CarveMe-GutMicrobes are identical for this species. The *B. breve* model generated with CarveMe-GutMicrobes shows very few false negatives (only four), resulting in high precision and specificity scores (Table S4). Together, the results from gene essentiality analysis further emphasize the value of integrating multiple resources to achieve robust metabolic reconstructions.

**Figure 4:**
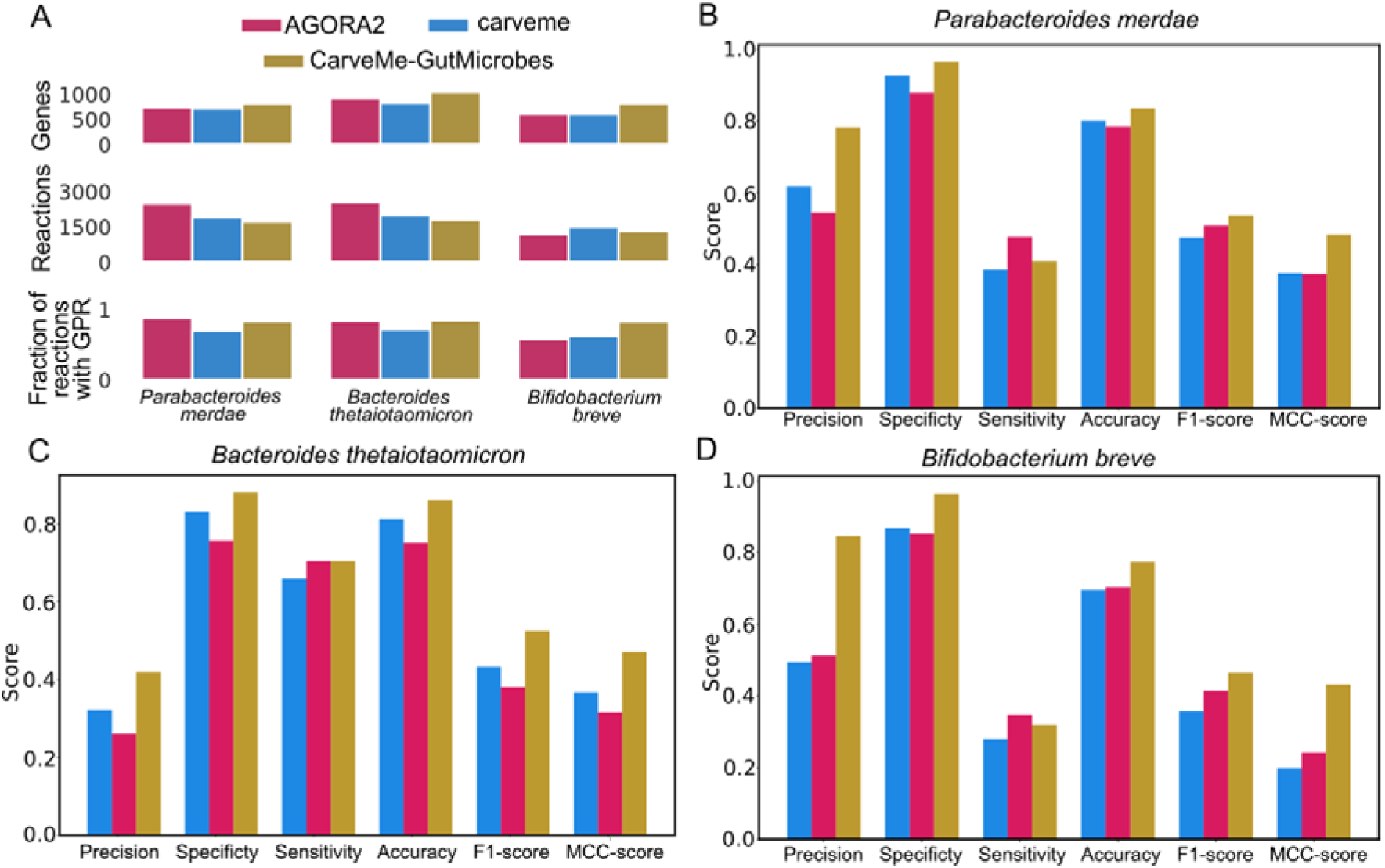
Results of the gene essentiality benchmark. (A) Summary of the number of genes, metabolites and reactions for each model drafted with CarveMe and CarveMe-GutMicrobes and those obtained from the AGORA2 collection. (B) Barplots reporting the benchmark results for *Parabacteroides merdae*, (C) *Bacteroides thetaiotaomicron*, and (D) *Bifidobacterium breve*.

Although CarveMe-GutMicrobes incorporates information from AGORA2, differences in sensitivity reflect their distinct reconstruction strategies. AGORA2 consists of manually curated, strain-specific fixed resource for metabolic models integrating extensive literature knowledge, enhancing detection of incompletely annotated pathways. In contrast, CarveMe-GutMicrobes relies on template-based reconstruction prioritizing completeness and growth feasibility, which can lead to underrepresentation of reactions explicitly curated in AGORA2.

##### Archaeal models

We evaluated the predictive performance of the archaeal GSMMs generated by CarveMe-GutMicrobes, CarveMe, and those available in the AGORA2 collection by comparing their ability to simulate metabolite uptake and secretion. This comparative analysis focused on four species, viz., *Methanobrevibacter ruminantium, Methanosphaera stadtmanae, Sulfolobus solfataricus, Methanobrevibacter smithii*, for which both curated models and experimental data were available. The constrained sample size reflects the limited representation of Archaea in AGORA2, which includes only seven archaeal reconstructions among 7032 total models. Experimental data were sourced from the NJC19 database (Lim *et al*., 2020), and simulation outcomes are summarized in Figure 5A. CarveMe-GutMicrobes and AGORA2 models exhibited comparable predictive accuracy for the available metabolite exchange profiles. CarveMe-GutMicrobes demonstrated improved accuracy over CarveMe in simulating key archaeal traits (Figure 5B), including methane and methanol metabolism, which are central to the ecological function of many archaeal taxa (Figure 5A). These findings highlight the current gap in archaeal GSMM resources and the potential of CarveMe-GutMicrobes to enable custom reconstructions and broader metabolic modelling of this underrepresented domain. Growth predictions in minimal media further support the enhanced functional coverage of CarveMe-GutMicrobes models.

**Figure 5:**
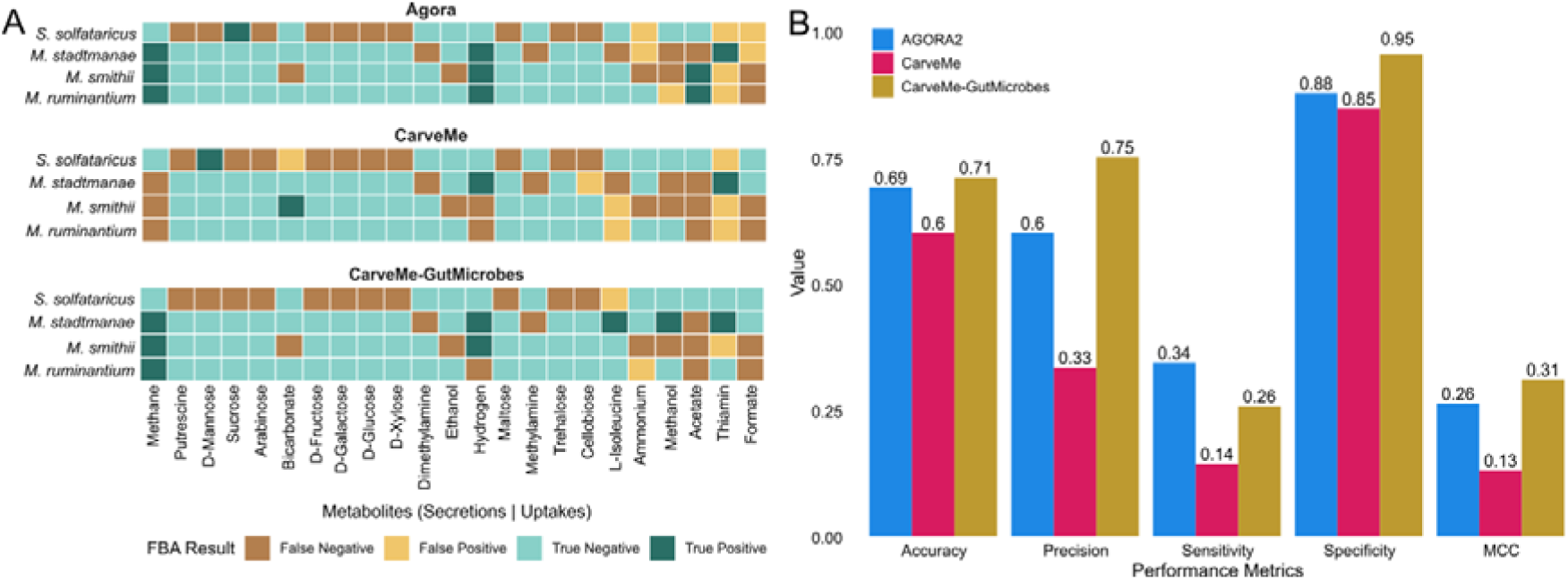
Comparison of the archaeal GSSMs constructed using CarveMe, CarveMe-GutMicrobes, and AGORA2 in reproducing the metabolic physiology data from the NJC19 database (Lim *et al*, 2020) (A) Summary of true-/false- positive/negative predictions for four archeal species for a range of secretion and uptake metabolites. (B) Barplot reporting the performance metrics (accuracy, precision, sensitivity, and specificity) for the GSMMs generated by CarveMe, CarveMe-GutMicrobes, and those obtained from AGORA2 collection.

### Diversity of Strain-Specific Metabolic Models

We reconstructed genome-scale metabolic models for all 30,691 genomes in the HumGut collection using both CarveMe and CarveMe-GutMicrobes. Models generated with CarveMe-GutMicrobes contained fewer reactions lacking supporting annotation evidence (i.e., GPR associations) (Figure 6A), indicating improved consistency with the underlying genomic data. To further assess the impact of the method on capturing strain-level metabolic variation, we selected all genomes from three representative species and compared pairwise differences using the Jaccard index. CarveMe-GutMicrobes models captured these inter-genome differences more accurately (Figure 6B, C, D), reflecting the distinct metabolic profiles of individual strains better than the original CarveMe models.

**Figure 6.**
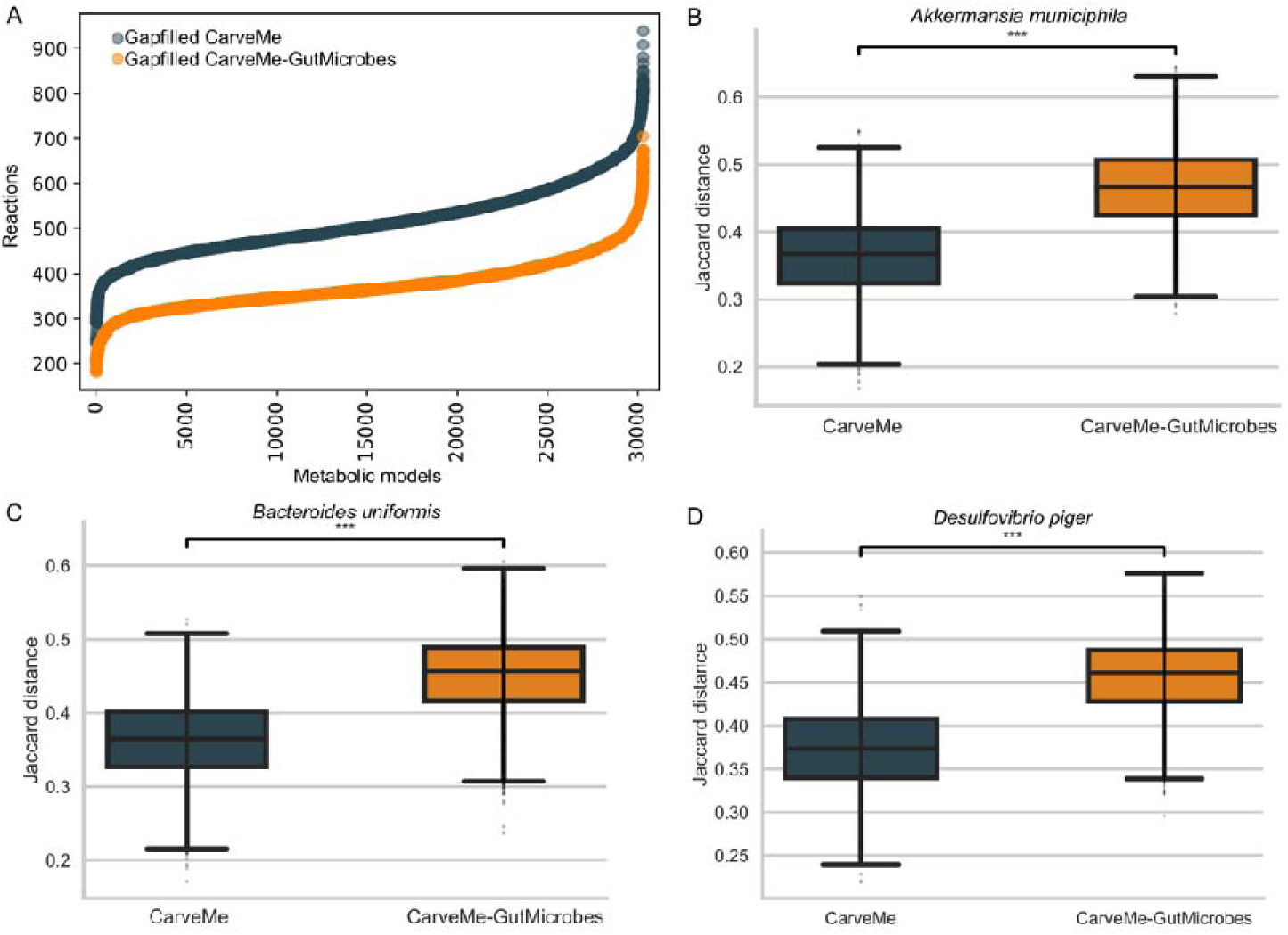
Comparison of metabolic models gapfilled reactions and inter-genome variability between CarveMe and CarveMe-GutMicrobes. (A) Distribution of reactions without annotation evidence (gapfilled) in genome-scale metabolic models reconstructed from HumGut genomes using CarveMe and CarveMe-GutMicrobes. (B–D) Pairwise Jaccard distances between models of the same species but different strains reconstructed with CarveMe and CarveMe-GutMicrobes for three representative gut organisms: (B) *Akkermansia muciniphila*, (C) *Bacteroides uniformis*, and (D) *Dorea piger*. Boxplots show the distribution of intrastrain metabolic dissimilarities. Statistical significance of differences between reconstruction methods was assessed using a paired Wilcoxon test.

### Computational performance of CarveMe-GutMicrobes

Benchmarks were conducted on the Cumulus partition of the CSD3 high-performance computing cluster (Intel Xeon Scalable CPUs, 3.5–6 GB RAM per core) of the University of Cambridge. Using a single CPU core, the mean reconstruction time per genome was 826.3 ± 717.9 seconds. Taxonomic information was consistently applied at the genus level. Runtime scaled with genome size and with the degree of representation of each taxon in the AGORA2 database and was additionally influenced by solver performance during network resolution, which varied according to genome annotation complexity.

The peak in the expected metabolic reconstruction time observed for Bacteroides spp. can be attributed to the high number of species within this genus (58 species). In total, there are 172 genomes attributed to such species, including 25 genomes from *Bacteroides fragilis* and 12 from *Bacteroides thetaiotaomicron*. The genomes are therefore distributed across multiple species, resulting in a high level of taxonomic diversity and increased computational complexity. For these taxa, a species-level taxonomic resolution would have been more appropriate; however, genus-level classification was adopted to ensure consistency with the taxonomic resolution used for the other genomes in the dataset.

In contrast, only 14 species of *Escherichia* are represented in the AGORA2 database, and 981 out of the 998 genomes representing this genus belong to *Escherichia coli*. Thus, most genomes correspond to different strains of the same species, leading to lower species-level diversity and explaining why the reconstruction time for *Escherichia* spp. is not comparably elevated despite the large number of genomes.

**Table 1:** Benchmarking results for genome-scale metabolic model reconstruction using CarveMeGut. For each gut-associated bacterial genome, the table reports genome size, reconstruction time (in seconds), and genus counts, defined as the number of occurrences of the corresponding genus in the AGORA2 database. Reconstruction time was measured using a single CPU core.

| Genome | Genome size | time (s) | Genus counts |
| --- | --- | --- | --- |
| *Prevotella copri* CB7 (DSM 18205) | 3.6 Mb | 396.48 | 68 |
| *Ruminococcus bromii* L2-63 | 2.2 Mb | 396.67 | 22 |
| *Roseburia intestinalis* L1-82 | 4.5 Mb | 434.18 | 19 |
| *Akkermansia muciniphila* ATCC BAA-835 | 2.9 Mb | 446.29 | 4 |
| *Lactobacillus rhamnosus* GG (ATCC53103) | 3.1 Mb | 449.4 | 82 |
| *Faecalibacterium prausnitzii* L2-6 | 3.2 Mb | 452.23 | 10 |
| *Bifidobacterium longum* DJO10A | 2.4 Mb | 665.65 | 124 |
| *Escherichia coli* K-12 substr. MG1655 | 4.6 Mb | 678.91 | 998 |
| *Bacteroides thetaiotaomicron* VPI-5482 | 6.3 Mb | 2,067.64 | 172 |
| *Bacteroides fragilis* NCTC 9343 | 5.2 Mb | 2,275.43 | 172 |

## Conclusion

We introduced CarveMe-GutMicrobes, an automated pipeline for the reconstruction of GSMMs tailored to human gut-associated microbes. Built upon a top-down reconstruction framework, CarveMe-GutMicrobes integrates gene-protein-reaction (GPR) associations from multiple community-curated resources, including AGORA2, UniRef90, TCDB, and BiGG, thereby aligning with the collective efforts of the COBRA modelling community. Comprehensive benchmarking against experimental phenotype data revealed that CarveMe-GutMicrobes models consistently match or outperform reconstructions generated using modelSEED, gapseq, and the original CarveMe pipeline, particularly in their ability to reproduce growth and metabolite secretion profiles. These findings underscore the enhanced predictive power and coverage of CarveMe-GutMicrobes models.

In addition to the modelling framework, our study also brings a latest experimental dataset on gene essentiality for prevalent gut bacteria. Given the importance of the availability of experimental datasets for building accurate models, esp. For non-model gut species, this study represents an important advance in metabolic modelling of human gut bacteria.

We therefore anticipate that CarveMe-GutMicrobes will serve as a valuable resource for linking genotype to phenotype in gut microbial ecosystems, and for accelerating the development of mechanistic insights into microbial community function.

## Data availability statement

All scripts can be accessed at https://github.com/arianccbasile/carvemegut, and the complete dataset is publicly available at https://zenodo.org/records/14882984.

## Supplementary materials

The following supplementary tables provide additional data and analyses supporting the main findings of this study:

- **Table S1**: Conversion of metabolic model IDs from BiGG identifiers to extended gene and metabolite names used throughout the analysis
- **Table S2**: List of essential genes identified in *Parabacteroides merdae*
- **Table S3**: Essential gene predictions for *Bacteroides thetaiotaomicron*
- **Table S4**: Essential genes identified in *Bifidobacterium breve*

## Acknowledgments

This project has received funding from the European Research Council (ERC) under the European Union’s Horizon 2020 research and innovation programme (project ModEM, grant no. 866028).

## Author contributions

AB: Conceptualization, Methodology, Software, Validation, Investigation, Data curation, Writing – original draft, Writing – review & editing, Visualization; IR: Gene essentiality data collection, Writing – review & editing; AM: Methodology, Validation, Visualization, Writing – review & editing; FZ: Methodology, Visualization, Writing – review & editing; SK: Methodology, Writing – review & editing; KRP: Conceptualization, Supervision, Project administration, Funding acquisition, Writing – original draft, Writing – review & editing.

